# Risk factor profile of different clinical manifestations of cerebral small vessel disease - data from SHEF-CSVD Study

**DOI:** 10.1101/167221

**Authors:** J. Staszewski, E. Skrobowska, R. Piusińska-Macoch, B. Brodacki, A. Stępień

**Affiliations:** Clinic of Neurology, Institute of Medicine, Szaserow 128, 04-141 Warsaw (POLAND); Department of Radiology, Military Institute of Medicine, Szaserow 128, 04-141 Warsaw (POLAND)

**Keywords:** cerebral small vessel disease, risk factors, lacunar stroke, vascular dementia, vascular parkinsonism

## Abstract

**BACKGROUND:** Little is known of the mechanisms of cerebral small vessel disease (CSVD). Both atherosclerosis or non-atherosclerotic diffuse arteriopathy are involved.

**METHODS:** A single-center, prospective, case-control study was performed in consecutive patients with different CSVD manifestations. The study group consisted of 205 patients: 52 with lacunar stroke (LS), 20 with subcortical hemorrhagic stroke (HS), 50 with vascular dementia (VaD), 28 with vascular parkinsonism (VaP) and 55 controls (CG) free of cerebrovascular disease but with high vascular risk.

**RESULTS:** Patients with CSVD had significantly higher prevalence of vascular risk factors including hypertension, diabetes mellitus, polymetabolic syndrome and chronic kidney disease. Patients with CSVD had also significantly higher fasting blood glucose, homocysteine, fibrinogen, systolic blood pressure, IMT values and lower eGFR, albumin and HDL levels. After adjustment for age and sex, low eGFR, albumin and high levels of uric acid and fibrinogen were associated with all CSVD groups, elevated fasting glucose was related to LS and HS. In the multivariate analysiss, the independent predictors for CSVD were female sex, low albumin, high fibrinogen, fasting glucose and uric acid. Patients with LS had significantly higher IMT values comparing to other CSVD groups, patients with VaP had a trend towards higher homocysteine levels.

**CONCLUSION:** Risk factor profile for CSVD as a whole differs from subjects with proatherogenic profile without history of cerebrovascular disease. Our results support the concept that CSVD is not homogeneous, and that unique risk factors profiles exist for different clinical manifestations of the disease.

## Introduction

Cerebral small vessel disease (CSVD) is one of the most important and common microangiopathy. It includes lacunar infarcts, intracerebral hemorrhages, white matter lesions (WMLs) or microbleeds. These lesions are frequent and often asymptomatic findings on CT and MRI scans of elderly people [1]. CSVD can cause several different types of distinct or overlapping clinical presentations: recurrent lacunar strokes (LS), deep haemorrhagic strokes (HS), vascular dementia (VaD) and vascular parkinsonism (VaP). Although CSVD is considered to result from cerebral arteriolar occlusive disease, classical cardiovascular risk factors are not consistently common in patients with CSVD. These findings are challenging the traditional view that classical risk factors play a role in CSVD genesis and indicate that pathophysiology of CSVD may be independent from that of atherosclerotic large artery disease [2]. It is also speculated that the exact mechanisms of distinct clinical CSVD manifestations differ and they may be attributable to either burden or control of traditional vascular risk factors but also are influenced by other hemodynamic or inflammatory factors [3].

The strength of association of vascular risk factors to different clinical spectrum of CSVD is difficult to ascertain because vascular lesions may accompany neurodegenerative mechanisms especially in patients with chronic CSVD manifestations: VaD or VaP. Due to lack of effective casual treatment, the identification of CSVD-specific modifiable risk factors is of increased importance for secondary prevention of brain lesions [4]. Although asymptomatic radiological CSVD markers f.e. WMLs or lacunes are frequently found in patients with coronary or peripheral artery disease, the comparison of risk factors profiles between patients with different manifestations of CSVD and patients with high vascular risk without cerebrovascular disease has not been reported so far. If atherosclerosis were important in CSVD as a whole or in one particular subtype, one would expect the risk factor profile to be similar or even aggravated to that of large vessel disease. Considering the wide spectrum of radiological and clinical picture of CSVD, we hypothesized that associated atherothrombotic risk factors differ between patients with CSVD and subjects without cerebrovascular disease but with high vascular risk.

In this single-center, prospective, case-control study, we compared prevalence of traditional risk factor profiles between patients with different CSVD manifestations and controls with high atherothrombotic risk free of clinical and radiological markers of CSVD.

## Materials and Methods

The present investigation is nested in the of SHEF-CSVD Study (Significance of HEmodynamic and hemostatic Factors in the course of different manifestations of Cerebral Small Vessel Disease) [5]. The study group consisted of 150 consecutive patients: with first-ever recent LS (n=52) or deep HS (n=20), VaP (n=28) and VaD (n=50) and 55 controls (CG). The patients were recruited from Neurological Outpatient Department and General Cardiology Outpatient Department and prospectively enrolled in the study between December 2011 and June 2014. The study protocol and methods have been thoroughly described elsewhere [5]. In brief, SVD group consisted of consecutive patients with a first ever recent LS or HS or newly diagnosed VaD and VaP presumed to be caused by SVD with evidence of typical findings on neuroimaging (MRI), who were independent (total Barthel Index >80 points) and had no severe dementia (MMSE≥12 points) [6]. The patients were diagnosed according to typical radiological and clinical picture: LS - according to the OCSP Criteria; VaD and VaP after exclusion of other neurodegenerative conditions with the use of clinical tools easily applied in clinical practice: Hurtig criteria or NINDS-AIREN criteria with Modified Hachinski Ischemic Scale≥ 7 points, respectively [7, 8, 9]. Patients with recurrent LS or strategic single-infarct dementia or with post-stroke VaD or VaP were excluded. Mean time from first symptoms to enrollment was 23,2±10 months in VaD and 25±10 months in VaP (p=0,5).

The control group consisted of patients without history of cerebrovascular disease and with high cardiovascular risk assessed according to the European Society of Cardiology and the European Atherosclerosis Society Guidelines (2011) [10]. High risk was recognized in patients with: documented cardiovascular disease (CVD) – previous myocardial infarction (MI), acute coronary syndrome, coronary revascularization/bypass graft or peripheral arterial disease (PAD); diabetes (type 2 or type 1 diabetes with target organ damage e.g. microalbuminuria); moderate to severe chronic kidney disease (CKD; glomerular filtration rate (GFR) < 60 ml/min/1.73 m2); or markedly elevated single risk factors such as familial dyslipidemias and severe hypertension (systolic blood pressure (SBP) ≥180 mm Hg and/or diastolic blood pressure (DBP) ≥110 mm Hg); or 10-year risk of total CVD ≥5% (estimated using the Systemic Coronary Risk Estimation (SCORE) risk assessment charts according to gender, smoking status, age, blood pressure and total cholesterol (TC)) [11].

To prevent confounding by hyperacute phase responses, all LS patients underwent study procedures at least 2 weeks (mean 19,4±4,1 days) after their index strokes. All participants were aged between 60 and 90 years. Patients with significant stenosis (>50%) of a major extracranial or intracranial artery, atrial fibrillation, non-SVD related WMLs (e.g. due to migraine, vasculitis, multiple sclerosis, CADASIL), life expectancy of less than 6 months, and MRI contraindications were excluded. Based on medical records, physical examination and comprehensive history available at baseline, we evaluated atherothrombotic risk factors, including tobacco use, diabetes, hyperlipidemia, hypertension, coronary artery disease (CAD) and peripheral vascular disease (PAD). To determine baseline blood pressure control we performed 24h ABPM using a portable non-invasive oscillometric and auscultatory device (Schiller MT-300). Hypertension was defined as persistent elevation of systolic blood pressure >140 mm Hg or diastolic blood pressure >90 mm Hg at least 1 week from stroke onset, or current treatment with antihypertensive drugs. Diabetes mellitus was defined as a previous diagnosis of type I or type II diabetes, or at least two random glucose readings of ≥200 mg/dL mmol/l or fasting blood glucose readings of ≥126 mg/dL. Hypercholesterolaemia was defined as a serum total cholesterol >200 mg/dL or current treatment with a statin. The following criteria were used to diagnose polymetabolic syndrome: waist circumference ≥102 cm in men or ≥88 cm in women; HDL < 40 mg/dL in men and <50 mg/dL in women or on drug treatment; elevated blood pressure ≥130 mmHg systolic or ≥85 mmHg diastolic or on drug treatment; elevated TG ≥150 mg/dL or on drug treatment; and elevated fasting glucose (FG) ≥100 mg/dL or on treatment for diabetes [12]. Coronary artery disease (CAD) was defined in patients with stable angina, prior myocardial infarctions (MIs), prior percutaneous revascularization, coronary artery bypass graft, angiographically proven coronary atherosclerosis, or reliable non-invasive evidence of myocardial ischemia [13]. Previous history of CAD and peripheral artery disease (PAD) was recorded based on clinical history and documented investigations. Assessments of carotid intima-media thickness (IMT) were performed according to previously validated criteria by colour-flow B-mode Doppler ultrasonography by a same experienced sonographer. The IMT was defined as the distance between the leading edge of the lumen-intima echo and the leading edge of the media-adventitia echo [14]. At least three measurements were taken over a 1-cm length of far wall of each CCA segment, and these measurements on both sides were averaged to obtain the mean IMT. For all participants, serum total cholesterol (TC), HDL, LDL, triglycerides (TG), fasting glucose, HbA1c, homocysteine levels were assessed. All studied patients had MRI examination with assessment of white matter lesions (WMLs) in Fazekas scale before entering the study. Grade 2 (n=83; 55,3%) or 3 (n=45;30%) WMLs were present in 80,3% patients with CSVD. Controls were included only in case of normal MRI scans (Grade 0).

This study complied with the Declaration of Helsinki. All participants from both groups signed an informed consent form. This study was approved by the local Medical Ethics Committee and was supported by the Polish Ministry of Science and Higher Education as a research project of the Military Institute of Medicine (Warsaw, Poland, study number N N402 473840).

## Statistical analysis

Categorical data were presented as frequencies and compared using Chi-square, factorial logistic regression, or Fisher’s exact test, where appropriate. Continuous data were reported as means±SD and compared using paired t tests, non-normal data were analyzed using non parametric tests. Difference between all groups was assessed by using ANOVA method. Logistic regression adjusted for age and sex was used to calculate odds ratios (ORs) and 95% confidence intervals (CI) and to assess the strength of association between studied variables and CSVD groups and CG. For continous variables the ORs per 1-SD increase was used. Statistical models were built sequentially: all variables with a *p* value <0.1 in univariate testing were entered into adjusted multivariate logistic regression models consisting of categorical data and continuous data. Final model (Model 2) was built using backward stepwise regression analyses. A probability value of p<0.05 was considered significant. All data are presented as mean±SD values. All analyses were performed using Statistica 12 software (StatSoft Inc, USA).

## Results

### Prevalence of vascular risk factors among patients with CSVD and controls

We assessed statistical significance of the risk factors between patients with CSVD and controls (Table 1). In the univariate analyses, mean age, sex distribution, frequency of smoking and hyperlipidemia, PAD and obesity were similar in CSVD and CG although patients with CSVD more often had polymetabolic syndrome, were diabetic, had elevated fasting glucose and HbA1c, had more frequent hypertension with increased 24h SBP, CKD and less often had recognized CAD compared to CG. Patients with CSVD had also significantly higher levels of homocysteine and increased IMT values but lower HDL. In CSVD subjects without diabetes or CKD, mean HbA1c and fasting glucose were also significantly elevated and eGFR was decreased compared with CG (respectively, 5,8±0,31 vs 5,6±0,31%, p=0,003; 98,4±10,38 vs 93,2±11,88 mg/dL, p=0,02; 80,4±23,22 vs 99,1±20,71 ml/min, p<0,01). There was no difference in frequency of bad control of glycaemia (HbA1c>7,5%) between diabetic patients from CG and CSVD (8,3% vs 20%; p=0,18), HS (5%,p=0,3), VaP (21%, p=0,2) and VaD (13%, p=0,6) but it was more frequent in patients with LS (31,23%, p=0,03). Compared with CG, after adjustment for age and sex, patients with LS were more likely to have hypertension with elevated SBP, diabetes, polymetabolic syndrome, CKD, increased BMI and FG, patients with HS were more frequently active smokers, had CKD and were less likely to have hyperlipidemia (Fig.1). There were no differences between CG and VaD and VaP in terms of frequency of hypertension, smoking, hyperlipidemia, and obesity. There was however a trend toward higher prevalence of diabetes, polymetabolic syndrome in patients with VaD. All CSVD subgroups had less frequent CAD, lower eGFR, albumin levels, elevated uric acid, fibrinogen than controls. In the multivariate analysis controlled for all significant varaibles from univariate analysis, the independent factors associated with CSVD group were female sex, low albumin, high levels of fibrinogen, plasma glucose and uric acid (Table 2). There was also a trend towards higher risk in patients with low eGFR and polymetabolic syndrome. The best adjusted statistical model consisted of IMT, albumin and fibrinogen. Low albumin and high fibrinogen levels were risk factors for all CSVD subgroups, female sex was associated with LS and VaD, high blood glucose and uric acid were independently associated with LS only.

**Table 1.** Demographics and laboratory data of patients with CSVD and controls.

|  | CG<br>(n=55) | CSVD<br>(n=150) | LS (n=52) | HS (n=20) | VaP (n=28) | VaD (n=50) | p<sup>#</sup> |
| --- | --- | --- | --- | --- | --- | --- | --- |
| Age (y) | 72 (5,9) | 72,4 (8,4) | 69,9 (8,7) | 74,1 (10,4) | 72,3 (6,24) | 74,4 (7,9)** | ,04 |
| Female sex (%) | 25 (45,5) | 76 (50) | 19 (37) | 10 (50) | 10 (36) | 37 (74)** | ,01 |
| Hypertension (%) | 43 (78,2) | 132 (88)* | 49 (94)* | 19 (95)* | 20 (71,4) | 44 (88) | ,01 |
| MAP (mmHg) | 90,64 (9,8) | 94,98 (13,05) | 95,02 (12,9) | 99,77 (19,4)* | 91,24 (11,14) | 95,24 (11,1) | ,46 |
| SBP (mmHg) | 125,3 (18) | 133,9 (17,2)** | 136,5 (18,3)** | 137,1 (21,8)* | 128,8 (13) | 132,9 (15,3)* | ,33 |
| DBP (mmHg) | 74,8 (8,2) | 75,53 (12,5) | 74,24 (11,7) | 81,1 (19,7) | 72,2 (10,3) | 77 (10) | ,32 |
| CAD (%) | 22 (40) | 29 (19)* | 10 (19)* | 3 (15) | 3 (11) | 13 (26) | ,39 |
| Diabetes mellitus (%) | 20 (37) | 81 (54)* | 29 (56) | 10 (50) | 14 (50) | 28 (56) | ,9 |
| HbA1c % | 5,9 (0,6) | 6,3 (1)** | 6,57 (1,22)** | 5,9 (0,4) | 6,3 (1) * | 6,2 (0,9)* | ,1 |
| FBG (mg/dL) | 103,1 (20) | 123,2 (44,9)** | 132 (51,47)** | 134,2 (44,6)** | 113 (32,7) | 115 (41,7)* | ,12 |
| Current smoking (%) | 15 (27,3) | 49 (32,7) | 18 (34,6) | 9 (45) | 11 (39,3) | 11 (22) | ,2 |
| Hyperlipidemia (%) | 43 (78) | 107 (71,3) | 39 (75) | 11 (55)* | 20 (71,4) | 37 (74) | ,37 |
| LDL (mg/dL) | 114,6 (36,9) | 108,6 (36,5) | 109,28 (38,8) | 112,45 (28,7) | 107,4 (34) | 109,5 (39,4) | ,81 |
| HDL (mg/dL) | 56,5 (17,5) | 51,3 (16,7)* | 45,9 (9,9)** | 58,7 (20,3) | 54,9 (13,5) | 50,8 (21,4) | ,01 |
| TG (mg/dL) | 126 (142) | 126 (76,8) | 147,9 (110) | 109,5 (46) | 126,3 (54,1) | 111,8 (46,6) | ,05 |
| TC (mg/dL) | 192,2 (38,9) | 182,9(42,4) | 178,6 (42) | 185 (34) | 187,5 (43,1) | 188,5 (45,3) | ,51 |
| PAD (%) | 13 (23,6) | 27 (18) | 8 (15) | 4 (21) | 6 (21,4) | 4 (8) | ,01 |
| BMI | 26,4 (4,3) | 27 (5,3) | 28,5 (5,99)* | 26,4 (5,4) | 25,4 (3,6) | 26,7 (4,8) | ,07 |
| Obesity (BMI>30) | 11 (20) | 37 (24,7) | 18 (34,6) | 2 (10) | 5 (18) | 12 (24) | ,12 |
| PS (%) | 13 (23,6) | 63 (42)* | 25 (48)** | 8 (40) | 9 (32) | 21 (42) | ,5 |
| CKD (%) | 3 (5,5) | 25 (16)* | 10 (19)* | 5 (25)* | 5 (18)* | 5 (10)* | ,4 |
| eGFR (ml/min) | 99,9 (22,3) | 77,2 (24,6)** | 78,21 (27,87)** | 74,3 (19,7)** | 70,2 (22,2) ** | 79,6 (22,4)** | ,38 |
| Albumin (g/dL) | 4,5 (0,52) | 3,7 (0,7)** | 3,7 (0,7)* | 3,6 (0,6)* | 3,9 (0,6)* | 3,7 (0,9)* | ,01 |
| Fibrinogen (mg/dl) | 284,2 (70,2) | 352 (86,9)** | 349,7 (76,6)* | 359,9 (78,9)* | 343,4 (99)* | 357,7 (95)* | ,4 |
| Uric acid (mg/dl) | 4,8 (1,2) | 5,9 (1,9)** | 6,3 (2,4)* | 5,8 (1,8)* | 5,9 (1,4)* | 5,4 (1,6)* | ,2 |
| Vit.B12 (pg/ml) | 238,5 (90) | 232,3 (119) | 209,1 (112) | 224 (130)* | 299,4 (113,5)* | 223,9 (117) | ,03 |
| Homocysteine (mg/dl) | 13,2 (5,1) | 15,3 (6,7)* | 13,8 (4,6) | 16,9 (7,6)* | 17,5 (7,5)** | 15,1 (7,5) | ,1 |
| IMT (mm) | 0,9 (0,1) | 1 (0,1)** | 1,1 (0,2)** | 1 (0,1) | 1 (0,1) | 1,1 (0,2)* | ,01 |
| Antiplatelet treatment | 24 (43,6) | 95 (63) | 33 (62)<sup>&</sup> | 9 (45)<sup>&</sup> | 19 (67,9) | 37 (74) | ,1 |
| Statin treatment | 31 (57,4) | 92 (61) | 29 (55)<sup>&</sup> | 10 (50)<sup>&</sup> | 17 (60) | 36 (72) | ,11 |

Values are means (±SD) for continuously distributed data or numbers (%) for categorical data

**Figure 1:**
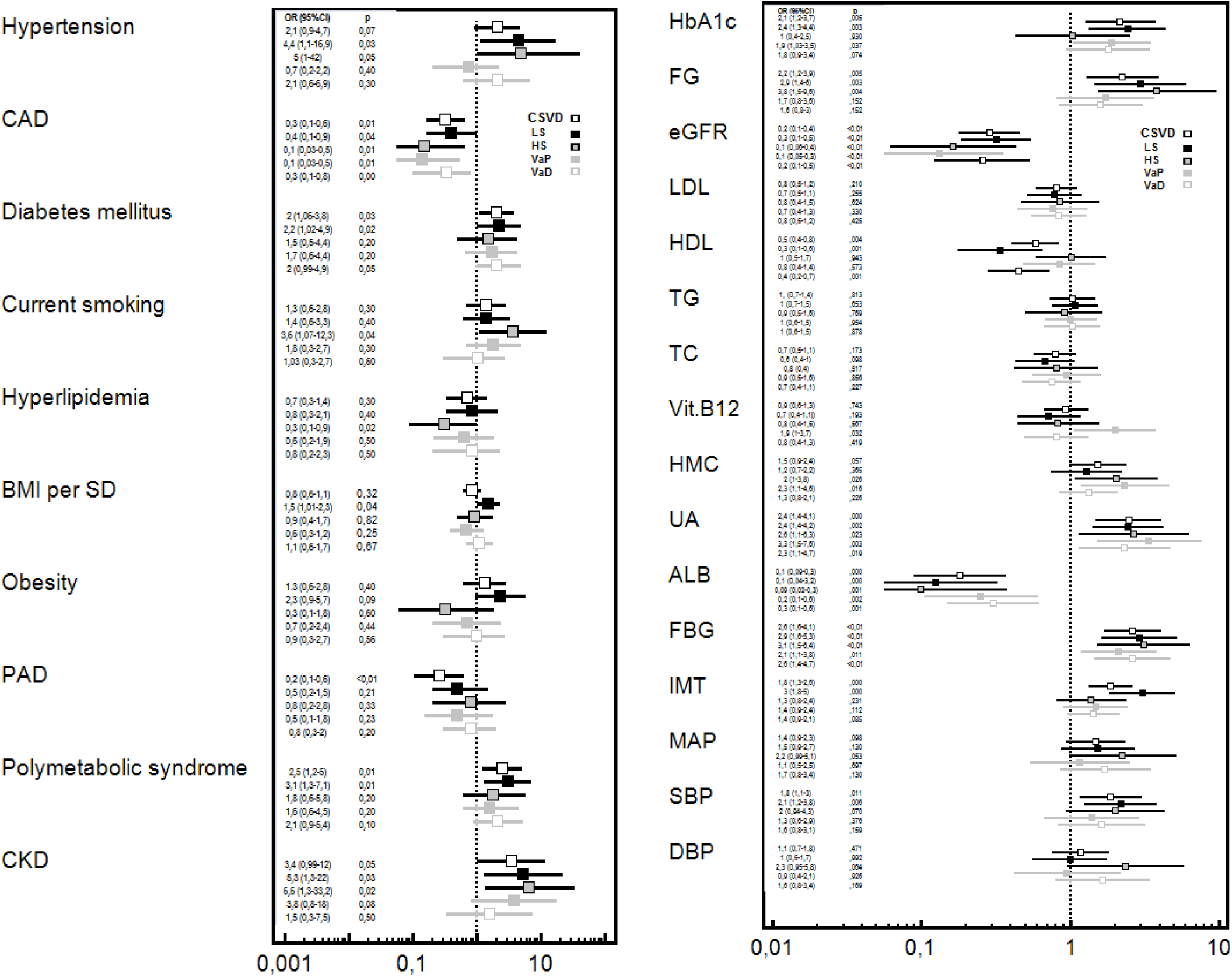
Comparison between risk factor profiles between CSVD subgroups controlling for age and sex. Data are expressed as OR (95% CI) CSVD – cerebral small vessel disease; CG – control group; LS – lacunar stroke; VaD – vascular dementia; VaP – vascular parkinsonism; HT-hypertension; CAD– coronary artery disease; DM–diabetes mellitus; CS–current smoking; HL–hyperlipidemia; BMI–body mass index; FG–fasting glucose; PAD – peripheral artery disease; PS–polymetabolic syndrome; IMT–intima media thickness; MAP–mean arterial pressure; SBP – systolic blood pressure; DBP–diastolic blood pressure, ALB – albumin

**Table 2.** Multivariate logistic regression analyses of variables associated with CSVD subgroups.

| Group | CSVD |  |  |  | LS |  |  |  | HS |  |  |  | VaP |  |  |  | VaD |  |  |  |
| --- | --- | --- | --- | --- | --- | --- | --- | --- | --- | --- | --- | --- | --- | --- | --- | --- | --- | --- | --- | --- |
| Variable | Model 1<br>OR<br>(95%CI) | p | Model 2<br>OR<br>(95%CI) | p | Model 1<br>OR<br>(95%CI) | p | Model 2<br>OR<br>(95%CI) | p | Model 1<br>OR<br>(95%CI) | P | Model 2<br>OR<br>(95%CI) | p | Model 1<br>OR<br>(95%CI) | P | Model 2<br>OR<br>(95%CI) | p | Model 1<br>OR<br>(95%CI) | P | Model 2<br>OR<br>(95%CI) | p |
| Age | 0,96 (0,8-1,07) | ,5 | 0,9 (0,7-1,02) | ,1 |  |  |  |  | 1,06 (0,8-1,3) | ,5 |  |  | 0,9 (0,8-1,1) | ,5 |  |  | 0,9 (0,8-1,1) | 0,7 |  |  |
| Sex (F) | 4,6 (1,1-18,5) | ,02 | 0,6 (0,1-4,7) | ,7 | 4,6 (1,1-18,5) | ,02 |  |  | 0,1 (0,1-10) | ,4 |  |  | 1,5 (0,2-10) | ,6 |  |  | 30 (3 - 59) | <0,01 | 10 (2,2-53) | <0,01 |
| Diabetes Mellitus | 0,64 (0,1-5,8) | ,6 |  |  | 0,1 (0,01-1,08) | ,06 |  |  |  |  |  |  |  |  |  |  |  |  |  |  |
| FBG | 3,5 (1,03-12) | ,04 | 8,3 (0,96-72) | ,05 | 3,5 (1,04-12,1) | ,04 | 12,5 (1,8-83) | <0,01 |  |  |  |  |  |  |  |  | 0,5 (0,1-2,8) |  |  |  |
| HbA1c |  |  |  |  |  |  |  |  |  |  |  |  | 1,6 (0,5-4,7) | ,3 |  |  | 0,2 (0,1-3,1) | ,2 |  |  |
| Hypertension | 1,1 (0,1-7) | ,9 | 0,6 (0,1-5) | ,6 |  |  |  |  | 0,2 (0,1-11) | ,4 |  |  |  |  |  |  |  |  |  |  |
| SBP | 1,17 (0,5-2,5) | ,6 |  |  |  |  |  |  | 2,1 (0,4-9,6) | ,3 |  |  |  |  |  |  |  |  |  |  |
| CKD | 3,4 (0,04-258) | ,5 | 10,9 (0,1-82) | ,2 |  |  |  |  |  |  |  |  | 0,8 (0,1-16) | ,8 |  |  | 0,3 (0,1-66) | ,6 |  |  |
| eGFR | 0,5 (0,2-1,1) | ,09 | 0,4 (0,2-1,1) | ,09 | 0,5 (0,2-1,08) | ,08 |  |  | 0,5 (0,1-4,1) | ,5 |  |  | 0,25 (0,1-1,4) | ,1 | 0,2 (0,1-0,7) | ,1 | 0,3 (0,1-1,2) | ,09 |  |  |
| PS | 6,9 (0,7-62) | ,08 | 0,6 (0,1-5,8) | ,7 | 5,7 (0,8-40) | ,07 |  |  |  |  |  |  |  |  |  |  |  |  |  |  |
| IMT | 1,5 (0,8-2,9) | ,18 | 2,1 (1,1-4,5) | ,04 | 1,8 (0,9-3,4) | ,07 |  |  |  |  |  |  |  |  |  |  |  |  |  |  |
| Homocysteine | 0,95 (0,4-2) | ,9 |  |  |  |  |  |  | 4,7 (0,4-57) | ,2 | 4,9 (0,6-37,7) | ,12 | 1,4 (0,6-3,3) | ,4 |  |  | 0,5 (0,2-1,4) | ,2 |  |  |
| Albumin | 0,36 (0,1-0,9) | ,04 | 0,1 (0,1-0,9) | ,04 | 0,35 (0,1-0,9) | ,03 | 0,1 (0,1-0,5) | <0,01 | 0,2 (0,1-1,1) | ,07 | 0,15 (0,1-0,9) | ,04 | 0,3 (0,1-1,5) | ,1 | 0,28 (0,1-0,9) | ,04 | 0,27 (0,1-0,9) | ,03 | 0,3 (0,1-0,9) | ,04 |
| Fibrinogen | 3,8 (1,6-9,1) | <0,01 | 3,1 (1,2-7,9) | ,01 | 3,8 (1,7-8,5) | <0,01 | 3,7 (1,3-10,7) | ,01 | 8,7 (0,9-81) | ,05 | 6 (1,1-31) | ,03 | 3,3 (1,2-9,3) | ,01 | 2,9 (1,2-7,1) | ,01 | 3,4 (1,2-9,2) | ,01 | 3,5 (1,4-8,4) | <0,01 |
| HDL | 0,7 (0,3-1,5) | ,4 | 0,3 (0,1-1,3) | ,1 |  |  |  |  |  |  |  |  |  |  |  |  |  |  |  |  |
| Uric acid | 2,4 (1,02-12) | ,04 | 1,5 (0,5-4,5) | ,4 | 2,8 (1,2-6,6) | ,01 | 2,7 (0,9-8,4) | ,07 | 3,2 (0,8-13,5) | ,1 | 2,6 (0,8-9) | ,1 | 1,4 (0,4-5) | ,6 |  |  | 1,5 (0,5-5) | ,4 |  |  |
| Vit.B12 |  |  |  |  |  |  |  |  | 1,1 (0,1-6,5) | ,9 |  |  | 1,2 (0,3-4,6) | ,7 |  |  | 0,5 (0,2-1,4) | ,2 |  |  |

### Difference between CSVD subgroups

Patients with VaD were on average 4,4 years (±1,6) older than patients with LS (p=0,04), the prevalence of females was the highest in VaD patients comparing to other CSVD subgroups (p=0,01). There were also significant differences between CSVD subgroups in ANOVA test with regard to IMT, vit.B12 and HDL levels. After adjustment for age and sex, patients with LS had lower HDL (difference between means 12,5±4,2 mg/dl (p=0,02), increased IMT and were more frequently obese than HS patients, had more prevalent hypertension and lower HDL than VaP (difference between means 8,8±3,9 mg/dl, p=0,04) and had increased IMT comparing with VaD and VaP patients (Table 3). Patients with HS had more prevalent hypertension than VaP subjects. Patients with LS and VaD had lower vit.B12 levels (difference between means, respectively 90,2±30 pg/ml, p=0,02 and 75,5±30 pg/ml, p=0,05) than VaP patients. There was no significant difference in CSVD groups with regard to levels of homocysteine, LDL, TG, TC and control of blood pressure.

**Table 3.** Comparison of risk factors profile between CSVD subgroups controlled for age and sex.

| Variable | LS vs HS | VaP vs LS | VaD vs LS | VaP vs HS | VaD vs HS | VaP vs VaD |
| --- | --- | --- | --- | --- | --- | --- |
| **Hypertension** | 0,7 (0,7-8,6) | 0,16 (0,3-0,6)* | 0,52 (0,1-2,5) | 0,1 (0,1-0,99) | 0,36 (0,03-3,4) | 0,29 (0,07-1,12) |
| **MAP** | 0,6 (0,3-1,2) | 0, 7 (0,4-1,4) | 1, 1 (0, 6-2,1) | 0,5 (0,2-1,2) | 0,7 (0,3-1,5) | 0,4 (0,2-1,1) |
| **SBP** | 0,9 (0,4-1,8) | 0,5 (0,2-1,1) | 0,7 (0,4-1,3) | 0,5 (0,2-1,4) | 0,7 (0,3-1,7) | 0,5 (0,1-1,3) |
| **DBP** | 0,5 (0,2-1,06) | 0,9 (0,5-1,8) | 1,6 (0,8-3) | 0,5 (0,2-1,2) | 0,7 (0,3-1,5) | 0,5 (0,2-1,2) |
| **CAD** | 1,8 (0,4-8) | 0,45 (0,1-1,8) | 1,07 (0,37-3) | 0,74 (0,12-4,32) | 2,02 (0,47-8,5) | 0,37 (0,08-1,5) |
| **Diabetes Mellitus** | 1,1 (0,4-3,3) | 0,88 (0,3-2,2) | 1,19 (0,4-2,9) | 0,9 (0,2-2,9) | 1,6 (0,5-4,9) | 0,59 (0,2-1,6) |
| **HbA1c%** | 2,5 (0,9-7,4) | 0,87 (0,5-1,4) | 0,82 (0,53-1,26) | 1,7 (0,7-4,1) | 2,1 (0,73-6,1) | 0,97 (0,56-1,68) |
| **FG** | 0,9 (0,6-1,4) | 0,6 (0,3-1,1) | 0,7 (0,5-1,1) | 0,5 (0,2-1,05) ! | 0,7 (0,4-1,1) | 0,7 (0,3-1,4) |
| **Current smoking** | 0,3 (0,1-1,2) | 1,3 (0,5-3,5) | 0,89 (0,32-2,4) | 0,66 (0,19-2,2) | 0,36 (0,1-1,2) | 1,9 (0,6-5,8) |
| **Hyperlipidemia** | 2,1 (0,6-6,6) | 0,84 (0,29-2,4) | 1,3 (0,4-3,6) | 1,9 (0,5-6,6) | 2,7 (0,84-8,6) | 0,83 (0,2-2,6) |
| **LDL** | 0,89 (0,5-1,5) | 1,03 (0,6-1,6) | 1,1 (0,7-1,8) | 0,8 (0,4-1,6) | 1 (0,5-1,8) | 0,7 (0,4-1,3) |
| **HDL** | 0,4 (0,2-0,8)* | 2,9 (1,3-6,4)* | 0,8 (0,5-1,4) | 0,8 (0,4-1,6) | 0,5 (0,3-0,9)* | 2,03 (1,1-3,7) |
| **TG** | 1,6 (0,6-4) | 0,7 (0,4-1,4) | 0,6 (0,2-1,2) | 2,1 (0,5-8,6) | 1,4 (0,3-5,8) | 1,9 (0,6-6,1) |
| **TC** | 0,8 (0,4-1,4) | 1,3 (0,8-2,1) | 1,2 (0,7-1,9) | 1,1 (0,5-2) | 0,9 (0,5-1,7) | 1,05 (0,6-1,7) |
| **BMI** | 1,4 (0,8-2,4) | 0,5 (0,3-0,9)* | 0,7 (0,3-1,4) | 0,94 (0,82-1,08) | 1,1 (0,6-1,9) | 0,5 (0,3-1,1) |
| **Obesity** | 4,9 (1-24,9)* | 0,43 (0,13-1,3) | 0,48 (0,1-1,3) | 1,75 (0,29-10,4) | 2,4 (0,4-12,8) | 0,63 (0,17-2,3) |
| **PS** | 1,1 (0,3-3,4) | 0,57 (0,21-1,5) | 1,01 (0,4-2,4) | 0,6 (0,18-2,1) | 1,4 (0,4-4,4) | 0,58 (0,2-1,6) |
| **PAD** | 0,8 (0,2-2,8) | 0,5 (0,15-2,8) | 1,3 (0,4-3,8) | 0,6 (0,14-3) | 1,0 (0,2-3,6) | 0,66 (0,18-2,3) |
| **CKD** | 0,7 (0,2-2,6) | 0,7 (0,2-2,6) | 0,4 (0,1-1,5) | 0,6 (0,1-2,5) | 0,3 (0,1-1,5) | 1,9 (0,4-8,5) |
| **eGFR** | 1,03 (0,6-1,7) | 0,8 (0,4-1,3) | 1,2 (0,7-2) | 0,7 (0,3-1,6) | 1,3 (0,6-2,5) | 0,6 (0,3-1,1) |
| **Vit.B12** | 0,93 (0,5-1,6) | 2,1 (1,2-3,8)* | 1 (0,6-1,7) | 1,9 (0,99-3,6) | 0,8 (0,4-1,0) | 1,9 (1, 1-3,3) |
| **Homocysteine** | 0,5 (0,2-0,9)* | 1,9 (1,06-3,6)* | 1,3 (0,8-2,1) | 1,04 (0,6-1,7) | 0,9 (0,5-1,4) | 1,1 (0,7-1,8) |
| **IMT** | 2,3(1,1-4,5)* | 0,4 (0,2-0,8)* | 0,49 (0,3-0,8)* | 0,9 (0,4-2) | 1,1 (0,6-2) | 0,8 (0,4-1,4) |
| **Albumin** | 1,04 (0,5-1,9) | 1,4 (0,8-2,7) | 1,08 (0,6-1,7) | 1,6 (0,7-3,8) | 0,9 (0,5-1,8) | 1,3 (0,7-2,5) |
| **Uric acid** | 1,1 (0,7-1,9) | 0,8 (0,5-1,3) | 0,7 (0,4-1,1) | 0,98 (0,4-2,2) | 0,8 (0,4-1,7) | 1,2 (0,6-2,4) |
| **Fibrinogen** | 0,8 (0,4-1,6) | 0,9 (0,5-1,5) | 0,9 (0,5-1,5) | 0,7 (0,4-1,4) | 1,07 (0,6-1,9) | 0,8 (0,4-1,3) |

\*p<0,05
HT- hypertension; CAD–coronary artery disease; DM–diabetes mellitus; CS–current smoking; HH–hypercholesterolemia; BMI–body mass index; FG–fasting glucose;
PAD – peripheral artery disease; PS–polymetabolic syndrome; IMT–intima media thickness; MAP–mean arterial pressure; SBP – systolic blood pressure; DBP–diastolic blood pressure

## Discussion

Our study documented that compared to controls free of cerebrovascular disease but with otherwise high vascular risk, patients with CSVD had significantly higher prevalence of vascular risk factors including hypertension, diabetes mellitus, polymetabolic syndrome and CKD. Patients with CSVD had also significantly higher fasting blood glucose, homocysteine, fibrinogen, systolic blood pressure, IMT values and lower eGFR, albumin and HDL levels. After adjustment for age and sex, low eGFR, albumin and high levels of uric acid and fibrinogen were associated with all CSVD groups, while elevated fasting glucose was related to acute CSVD manifestations: LS and HS. After controlling for all vascular comorbidities, the independent predictors for CSVD were female sex, low albumin, high fibrinogen, fasting glucose and uric acid. There was a trend higher risk in patients with lower eGFR and polymetabolic syndrome. Risk factors for different SVD manifestations were similar but comparing with other CSVD groups, patients with LS had significantly higher IMT values, those with VaP had a trend towards higher homocysteine levels.

Our results are in line with previous studies. In a systematic review of 16 studies comparing risk factors between patients with different stroke etiology, hypertension and diabetes were more frequent in patients with lacunar than in large vessel strokes [15]. There was no association between smoking, past cerebrovascular incidents and hypercholesterolemia with any type of ischemic stroke. In the study of Khan et al patients with lacunar strokes more frequently had hypertension whereas smoking, hypercholesterolemia, myocardial infarction, peripheral vascular disease were more common in nonlacunar stroke [16]. As we recruited patients with marked WMLs (presumably related to lipohyalinosis) and without ultrasound markers of large vessel disease it is not surprising that the most important risk factors in CSVD group were diabetes and metabolic disturbances (decreased eGFR) and not hypercholesterolemia and MI which are strongly related to large vessel disease.

After adjustment for age and sex, we showed a trend toward elevated HbA1c and fasting glucose especially in patients with ischemic lesions, increased BMI in LS patients but no association with BMI or obesity in other CSVD groups. That suggests that patients who have abnormally glycosylated end products, as are present in diabetes, may have more lipohyalinosis, while abnormal central obesity predisposes to lacunes by a distinct mechanism. This is in line with some reports indicating that patients with lacunar infarcts caused by atheromatous plaque in penetrating artery had the highest prevalence of CAD and patients with LS due to lipohayalinosis had the highest prevalence of diabetes and lowest prevalence of asymptomatic cerebral atherosclerotic disease [17, 18]. Similarly in the crosssectional ARIC study of 1827 community-dwelling participants, incident lacunes related to lipohyalinosis were associated with diabetes and HbA1c while LDL cholesterol, hypertension and smoking were associated with lesions presumable caused by microatheroma [19].

Our results demonstrated that low albumin, high fibrinogen were independent risk factors in different clinical manifestations of CSVD but there were also important differences: female sex was an independent risk factor in LS and VaD only; elevated uric acid, diabetes, polymetabolic syndrome and higher IMT values were associated with LS. Diabetes was a significant risk factor for LS and VaD when controlling with age and sex, but it was not significant when controlling for other vascular risk factors. This is in line with community-based cross-sectional studies which failed to find an association between diabetes and WMH and in Helsinki Aging Brain study in which WMLs were associated with diabetes only in persons <75 years of age [20, 21]. Higher levels of fibrinogen were associated with each of CSVD group what may support the hypothesis that coagulation pathway contributes to the pathogenesis of SVD. This association was independent of age, other cardiovascular risk factors and type of SVD manifestations regardless they were subacute (LS, HS) or chronic (VaD or VaP). Fibrinogen is an inflammatory marker, and a marker of systemic hypercoagulability acting as an important factor in the coagulation cascade. It is also assumed to be a faithful marker of brain-blood-barrier dysfunction. Probably the influence of systemic factors on the endothelium results in increase in permeability of the barrier and initiate pathologic changes. High fibrinogen levels serve as nonspecific marker for inflammatory disease, but also might predispose to thrombosis, reduce blood flow, enhance atherogenesis and lead to increased risk of cerebrovascular diseases. In the ASPS study, higher serum fibrinogen levels were independently associated with both WMLs and lacunar lesions on MRI [22]. In patients with symptomatic CSVD, fibrinogen was correlated with the amount of leukoaraiosis in patients with lacunar strokes and VaD [23]. Also in a systematic review and meta-analysis, levels of fibrinogen were elevated in chronic lacunar stroke versus non-stroke patients [24].

Our study showed low albumin level in all CSVD groups compared with controls. This finding is in line with several studies which suggested an inverse association between serum albumin concentrations and stroke risk. [25]. The underlying pathophysiology, however, remains unclear. Albumin increases the plasma oncotic pressure, decreases red blood cell sedimentation and viscosity which might favor reperfusion and leads to a better microvascular circulation. It is also recognized as an important antioxidant and thus reduced albumin may be a marker of chronic systemic inflammation [26]. It has been hypothesized that strokes due to CSVD can be caused by blood–brain barrier derangement with leakage of serum albumin and other toxins [27]. The cerebrospinal fluid/plasma albumin ratio, which was not available in this study, may be helpful for addressing this issue.

Patients with VaP had higher homocysteine and HDL levels while higher BMI and hypertension were less prevalent than in LS group. Beside an increased IMT which is a marker of subclinical atherosclerosis, there was no significant difference between LS and VaD. There was a similar risk factors profile between VaP and VaD patients although the later had lower HDL and vit.B12 levels. These different risk factor associations with CSVD subtypes may suggest that concomitant neurodegenerative process play an important role in the VaD and VaP genesis. Elevated level of HC can result from a folate deficiency in healthy individuals and it can be aggravated in patients with CKD. In our cohort we found that HC was associated with CSVD after controlling for age and sex especially in patients with VaP and HS and marginally in VaD but it was not an independent risk factor after controlling for other risk factors. That positive association with the presence of WMLs is well documented especially in patients with cognitive impairment [28]. Similar to Rotterdam Scan Study study, we found that hiperhomocysteinemia, low eGFR, elevated PGL and uric acid were independent of age and sex risk factors for the all CSVD subgroups however only uric acid in LS and eGFR in LS and VaD patients papered to be independent of other risk factors [29]. These findings suggests similar mechanisms involved in LS and VaD, probably related to endothelial activation caused by direct excititoxic effect of homocysteine, uric acid or uremic metabolites which appear to play a role in either acute (LS) or chronic (VaD) CSVD manifestations. These results are in line with the Northern Manhattan Study and the Rotterdam Scan Study which found that subjects with reduced eGFR had a greater burden of WMLs volumes after controlling for other factors [30, 31]. There is mounting evidence that chronic kidney disease increases the risk of different cerebrovascular disease including HS and subclinical WMLs [32]. Several biologically plausible mechanisms activated by CKD could result in development of WMLs through oxidative stress, procoagulant pathways activation, inflammation, resulting in endothelial dysfunction and vascular damage [33].

Although smoking was an independent of age and sex risk factor for HS, we did not find a correlation with LS,VaD, VaP while majority of studies documented that it is a risk factor for WMLs [34]. We found that smoking, hypertension were independent of age risk factors for HS and there was a trend toward higher MAP, SBP and DBP compared to controls. These results are consistent with previous trials which found that hypertension, smoking, alcohol consumption, low cholesterol and antithrombotic and statin treatment are linked to subcortical hemorrhage [35, 36]. Findings on the association between cholesterol levels and CSVD are not consistent. LDL, total cholesterol and triglicerydes were not independent risk factors for any CSVD subgroups, only low HDL was significantly associated with VaD and LS when adjusted for age and sex. That was in line with the LADIS study results which documented that among 639 elderly subjects with some degree of WMLs on MRI, incident lacunes were associated with low HDL. Another studies found that lower total cholesterol was associated with CSVD and mid-life lower HDL level was associated with late-life WMLs [37, 38, 39]. Also in a population based cohorts, elevated triglycerides and decreasing LDL were associated with severity of all MRI markers of CSVD [40]. In majority of the studies, the causal relationship between biomarkers cannot be determined because majority of them had cross-sectional design. We cannot directly compare our results with others because studies on CSVD usually concentrated on LS patients only or asymptomatic patients with WMLs.

Little is known about role of biochemical markers in CSVD pathophysiology and also of the magnitude of effect of classical vascular risk factors in the disease, which makes our data the more important. The advantages of the current study also include the relatively well characterized and simultaneously studied patients with different CSVD manifestations, the use of MRI in controls which enabled us to exclude patients with silent radiological markers of CSVD. We also enrolled a well-phenotyped group of patients with rarely studied chronic VaP and VaD. On the other hand our study has some limitations. The major weakness is the potential for random error or selection bias because of the small number of patients and controls included. Although patients with VaP and VaD were included to the present study immediately after diagnosis but they were in an advanced stage of their disease therefore it remains unknown whether our results can be applied to less severely affected patients.

In summary, our study showed that the risk factor profile for CSVD as a whole differs from subjects with proatherogenic profile without history of cerebrovascular disease. Our results support the concept that CSVD is not homogeneous, and that unique risk factors profiles exist for different clinical manifestations of the disease. Although this observation requires replication to ensure validity, if validated, it lends support to the involvement of multiple or different pathways in the pathogenesis of LS, HS, VaD or VaP. It is important that close association of CSVD with vascular risk factors gives a chance for successful primary and secondary prevention. The identified high-risk patient groups should be subjected to aggressive management of the underlying diseases and more close follow-up.

## Conflict of Interest

The authors have no conflict of interest to report.

